# The impact of behavioural activity on the EEG power spectrum, its source localisation, and global functional connectivity in rats

**DOI:** 10.64898/2026.07.03.736278

**Authors:** Čestmír Vejmola, Stanislav Jiříček, Marcel Bochin, Vlastimil Koudelka, Tomáš Páleníček

**Affiliations:** Psychedelic Research Center, National Institute of Mental Health, Topolova 748, Klecany, 25067, Czechia; Third Faculty of Medicine, Charles University in Prague, Ruska 87, Prague, 10000, Czechia; Clinical Research Program, National Institute of Mental Health, Topolova 748, Klecany, 25067, Czechia; Department of Complex Systems, Institute of Computer Science of the Czech Academy of Sciences, Prague, Czechia; Department of Cybernetics, Faculty of Electrical Engineering, Czech Technical University in Prague, Prague, Czechia; Faculty of Science, Charles University, Albertov 6, Prague, Czechia

**Keywords:** EEG, ECoG, source-localisation, connectivity, theta, thalamus, behaviour

## Abstract

The behavioural activity of freely moving animals is a confounding factor that affects the recording, analysis, and final results of animal EEG experiments. Along with the lack of standardisation in animal in vivo electrophysiology experiments, this could lead to huge inconsistencies, especially in the analysis of centrally acting drugs. Therefore, the main aim of this paper is to investigate the effects of behavioural activity versus inactivity on the multichannel EEG in freely moving rats. In a large sample (n = 116) of waking recordings from 12 cortical electrodes (ECoG) in Wistar rats, we evaluated behavioural activity-related changes in the power spectrum, current source density, and power-based global functional connectivity (GFC) in a 3D rat brain model, according to the TOHOKU Rat Brain Atlas. The main findings were that behavioural activity induced 1) a robust power increase in 6-8 Hz, peaking at 7 Hz with maximum changes over the parietal and temporal cortex, 2) an increase in gamma power (30-80 Hz) across the whole brain, 3) a decrease in delta (1-4 Hz) and beta (12-30 Hz) power across the whole cortex. Changes were also localised in subcortical regions, particularly in the diencephalon/thalamus. The GFC analysis showed a similar pattern of power changes across the 6-8 Hz, delta, and beta bands; however, GFC in the gamma band decreased. Again, the GFC analysis revealed changes in connectivity within subcortical structures, primarily in the thalamus. None of the measures was affected in the alpha band (8-12 Hz). These findings emphasise behavioural state as a critical factor influencing EEG outcomes, with important implications for the standardisation and translational validity of preclinical neurophysiological studies.

## Introduction

Standards and guidelines for EEG recording and analysis in animal research are still lacking due to non-standardized electrode positions, recording parameters, electrode types, and other methodological differences. As a result, the translational validity of animal EEG to human EEG remains limited. Compared with human studies, where data are typically collected from electrodes covering the entire brain, most animal studies focus on a specific region and use fewer electrodes. Consequently, they fail to reveal global effects, further limiting the use of modern computational methods for source localisation, as in human research (1).

Other significant sources of variability are related to behavioural activity (BA) versus inactivity and to open- versus closed-eye conditions. While in humans the resting-state (RS) EEG is usually recorded under clearly defined conditions of open/closed eyes, this is not possible in freely moving, awake animals. In such cases, differences between BA and RS during active, fully awake states, as well as the uncontrolled eye state, can lead to significant confounds. These effects are inevitably amplified when centrally acting drugs are used, as they typically alter the locomotor activity of animals (e.g., stimulants vs. sedatives, serotonergics, antipsychotics).

Our group has previously introduced a 12-electrode cortical electroencephalography (ECoG) system developed at the National Institute of Mental Health (Klecany, Czech Republic) and evaluated its performance for electrical source imaging (ESI) in rats. Recent advances in data collection, as well as in the development and adaptation of methods for 3D brain mapping (2), source localisation, and functional connectivity in the rat model (1,3–5), have enabled us to extend the current understanding of electrophysiological changes associated with BA and provide more translationally valid results.

In our studies, we have collected an extensive dataset using the 12-electrode system, with electrodes placed over the frontal, parietal, and temporal cortices of both hemispheres, and a reference electrode over the olfactory bulbs. This placement, together with previous evaluations of source localisation and conductivity in rat head phantoms and forward models, enables the effective use of whole-brain inverse source localisation methods, including the localisation of thalamic components of event-related potentials (ERPs) (1,6–8).

In addition to electrode-based functional connectivity measures (4,5,9,10), we can now compute functional connectivity from source-localised data, such as power-based global functional connectivity (GFC). This metric, commonly used in fMRI research, represents the average connectivity of a brain area with the rest of the brain (11–13). The EEG band-limited power (BLP) used for GFC computation is a strong correlate of the BOLD signal in fMRI (14–16) and shows a similar connectivity structure to BOLD (17,18). Our recent EEG–fMRI integration results (19) underscore the importance of using both modalities, as connectivity structures, though sometimes spatially similar, can carry different functional meanings.

In our previous pharmaco-EEG studies, we observed a substantial behavioural influence on ECoG changes induced by ketamine, amphetamine, and several psychedelics (3–5,10,20). It has also been suggested that BA, particularly running, induces a pronounced peak in the theta band at 6–8 Hz (theta peak) and a concurrent spectral power decrease in the delta and beta bands (21) in rats. However, in that study, only four electrodes were used over the motor and parietal cortices, limiting the ability to localise the observed changes.

Taken together, these findings motivated us to implement systematic co-registration of BA and RS conditions in all subsequent experiments. The present study aims to determine how BA affects the EEG in freely moving rats. Importantly, unlike many previous animal EEG studies, which are limited by small sample sizes and high inter-individual variability, our work benefits from a large dataset of N = 116 ten-minute baseline recordings collected over an extended period without pharmacological manipulation. This dataset allows for a statistically robust evaluation of RS/BA-related changes in EEG power spectra, source localisation, and global functional connectivity.

## Materials and methods

### Animals

Experiments were conducted on adult, experimentally naïve male Wistar rats (Velaz s.r.o., Konárovice, Czech Republic). Each animal was used in a single experiment. Rats were acclimated to the animal facility for at least 1 week prior to surgery and housed in pairs in plastic cages under a 12-h light/dark cycle (lights on at 6:00 a.m.), with a temperature of 21–24 °C and a relative humidity of 60%. Standard diet (ST2 pellets) and water were available ad libitum. All experimental procedures were approved by the Expert Committee for Protection of Experimental Animals of the 3rd Faculty of Medicine, Charles University (Prague, Czech Republic) and conducted in accordance with the Czech Animal Protection Act and EU Council Directive 86/609/EU.

### Stereotactic surgery

Rats were anaesthetised with isoflurane (2.5–3%) and positioned in a stereotactic frame (Stoelting) using atraumatic ear bars. The scalp was shaved, disinfected (betadine), incised, and the periosteum removed. Coordinates for electrode implantation were taken from the Paxinos Rat Brain Atlas (22), referenced to bregma: frontal association cortex (F3/F4: A + 5.0, L ± 2.0 mm), primary motor cortex (C3/C4: A + 2.2, L ± 3.2 mm), medial parietal association cortex (P3/P4: A − 3.8, L ± 2.5 mm), lateral parietal association cortex (P5/P6: A − 4.5, L ± 4.5 mm), secondary auditory cortex (T3/T4: A − 3.6, L ± 7.2 mm), and temporal association cortex (T5/T6: A − 8.3, L ± 5.8 mm). Holes (0.5 mm) were drilled using a Meisinger micro-drill, and gold-plated electrodes (Mill-Max, 0.48 mm tail) were implanted so that the tips touched the dura. Twelve recording electrodes were implanted bilaterally, with a reference above the olfactory bulb and a ground electrode subcutaneously in the occipital region. Electrodes were secured with Dentalon® cement. Animals recovered without weight loss or infection and were housed individually to protect the implant. One day before recordings, a connector was attached under brief isoflurane anaesthesia.

### Data collection

Data were collected over a series of pharmaco-electroencephalography (pharmaco-EEG) experiments, in which each animal underwent a baseline recording before any pharmacological intervention. Over the course of more than two years, this resulted in over 132 baseline EEG recordings from freely moving rats. Some of these datasets were excluded from subsequent analyses following the quality control procedure described below. Recordings took place seven days after surgery, during the lights-on phase of the light/dark cycle. Approximately 10 minutes prior to data acquisition, animals were connected to the EEG apparatus in their home cages to habituate to the setup. The recording session began with 10 minutes of baseline (no pharmacological manipulation), followed by drug administration in the original experiments (data not presented here).

Throughout the sessions, rats were allowed to move freely in their home cages while tethered via a lightweight cable to the amplifier. EEG was acquired with the BrainScope system (M&I, Prague, Czech Republic) at 16-bit resolution (7.63 nV/bit; ∼130 bit/μV), ±500 μV dynamic range, and 1 kHz sampling rate. Behavioural states were continuously monitored by a trained observer, who marked events in real time using predefined keyboard keys. Behavioural activity (BA) epochs included grooming, sniffing, rearing, and locomotion; resting state-like (RS) epochs represented periods of immobility. Artefact markers were used when the animal had to be handled (e.g., to untangle cables); these intervals were excluded from all further analyses.

### EEG signal preprocessing

Data were subsequently segmented into BA and RS epochs for three primary analyses: A) spectral power analysis at the electrode level, B) source-space power (POW) analysis, and C) global functional connectivity (GFC), performed across standard frequency bands: delta (δ, 1–4 Hz), theta (ϑ, 4–8 Hz), alpha (α, 8–12 Hz), beta (β, 12–25 Hz), high-beta (h-β, 25–30 Hz), gamma (γ, 30–40 Hz) and high gamma (h-γ, 55–100 Hz), as well as a narrow 6–8 Hz theta peak associated with running behaviour (23). The chosen band definitions were based on methodological standards proposed for rodent EEG and widely used in translational research, aiming to align preclinical frequency ranges with those applied in human EEG studies. Two additional analyses were conducted for the 6–8 Hz band: D) seed-based functional connectivity (SFC), and E) cross-frequency coupling (CFC).

Signal preprocessing was performed using BrainVision® Analyzer 2.1. Signals were re-referenced to the average reference (AVG), band-pass filtered between 0.5 and 100 Hz (8° slope), and notch filtered at 50 Hz. All recordings were visually inspected by an experienced EEG specialist to remove gross artefacts, including contaminated epochs and malfunctioning electrodes. Behavioural segmentation was based on the event markers described above, yielding quasi-continuous BA and RS intervals. Only datasets containing at least one minute of clean EEG in each condition were retained. Of the original 132 recordings, 116 met this criterion; therefore, the signal from 116 rats was analysed. For source localisation and connectivity analyses, data were exported from BrainVision® Analyzer using an in-house MATLAB (MathWorks, Natick, MA, USA) script.

### Spectral power analysis

Preprocessed data were segmented into 2-second epochs with 75% overlap. Fast Fourier Transform (FFT) with the following parameters: 0.5 Hz resolution, voltage output, non-complex, half-spectrum. A Hanning window (length = 10% of the segment, periodic, variance-corrected) was applied, and segments were zero-padded prior to transformation. FFT was applied to each epoch, and mean spectral power was calculated for each subject, each electrode, and each condition (BA and resting state, RS). Differences in spectral power between BA and RS conditions were statistically evaluated using a cluster-based permutation test (24) implemented in the FieldTrip toolbox (25). To identify the neural generators underlying BA–RS power differences, we applied a source localisation approach developed by our group (1). A homogeneous isotropic rat brain model (7) with realistic geometry and co-registered 12-electrode positions (Fig. 1C) was used to compute the forward model via the Finite Element Method (FEM) implemented in FieldTrip (26). The brain volume was sampled using a 1 mm isotropic grid. EEG time series were band-pass filtered to the frequency ranges defined above, segmented into 2-second epochs, and projected onto the grid of equivalent dipoles (voxels) using the eLORETA algorithm (27). For each voxel, a single representative time series was derived as the first principal component of the three orthogonal dipole moment time series. The mean power across all segments was then calculated for each voxel in each subject. Group-level differences in source-space power (POW) between BA and RS were tested separately for each frequency band using a nonparametric, cluster-corrected dependent t-test (24) (cluster α = 0.05; p < 0.05, two-sided, corrected). Significant t-values were mapped onto the TOHOKU Rat Brain Atlas™ (28) to enable identification and interpretation of functional brain regions exhibiting significant changes in EEG power between conditions.

**Figure 1.**
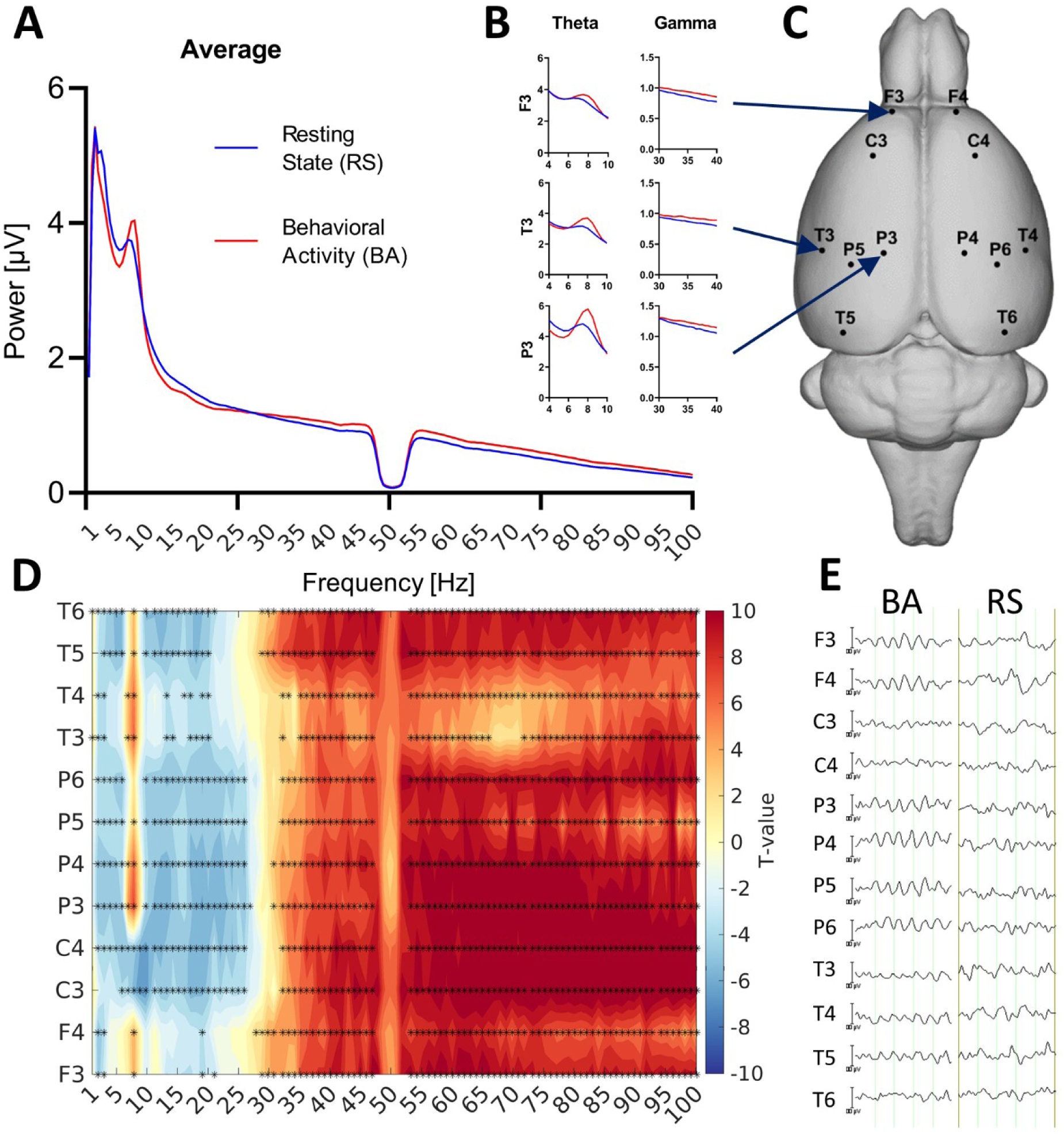
Spectral characteristics of EEG during behavioural activity (BA) and resting state (RS) in rats. **A)** Grand-average EEG power spectra (1–100 Hz) across all 12 electrodes and 116 rats during BA (red) and RS (blue). A prominent theta peak and global gamma power increase during BA are evident. **B)** Power spectra from selected electrodes (F3 – frontal, C3 – central, P5 – parietal) showing changes in theta (4–10 Hz) and gamma (30–40 Hz) bands. **C)** 3D rendering of the rat brain showing positions of all 12 epidural ECoG electrodes. **D)** T-statistic map showing significant spectral power differences (BA vs RS) across all electrodes and frequencies (1–100 Hz). Warmer colours indicate an increase in power during BA; cooler colours indicate a decrease in power. Black dots mark significant differences (p < 0.05, cluster-based permutation test). **E)** Representative 1-second segments of raw EEG traces from all 12 electrodes during BA (left) and RS (right), illustrating characteristic waveform differences.

### Global functional connectivity (GFC) analysis

Global functional connectivity (GFC) was assessed for each predefined frequency band. Source-projected EEG data were first downsampled to 250 Hz. For each dipole, seed-based connectivity was computed using the orthogonalized amplitude envelope correlation (oAEC) method (18). In this approach, the time series of each tested dipole was orthogonalized with respect to the seed dipole time series to minimise signal leakage. The analytic signal envelopes of both the orthogonalized and seed time series were then obtained as the absolute values of their Hilbert transforms. Connectivity between the seed and each tested dipole was quantified as the absolute value of the Spearman correlation between their envelopes. To reduce computational load, the number of tested dipoles per seed was downsampled by a factor of eight. For each recording, a subset of tested dipoles was randomly selected prior to analysis. For each seed dipole, GFC was defined as the median connectivity across the selected tested dipoles, yielding a single GFC map for each recording and frequency band. At the group level, differences in GFC between BA and RS conditions were evaluated using a non-parametric, cluster-based permutation test (24,25) applied to the GFC maps for each frequency band. Significant t-values were interpolated onto the TOHOKU Rat Brain Atlas™ (28), enabling anatomical interpretation of functional connectivity changes associated with behavioural activity.

### Seed-based functional connectivity (SFC)

To further focus on the narrow-band theta peak (6–8 Hz) frequency component, a seed-based functional connectivity (SFC) analysis was performed. Seeds were defined in regions showing the strongest source-level power (POW) increase at the theta peak. Based on the group-level t-maps, the right primary visual cortex (V1M) and the bilateral hippocampal formation were identified as major hubs, since these regions exhibited the most pronounced BA–RS differences (V1M t ≈ 5.7; hippocampus t ≈ 5.7 bilaterally). Both seeds were therefore selected for SFC analysis to capture the complementary roles of sensory and mnemonic circuits. The connectivity of each seed with all other dipoles was calculated using the orthogonalized amplitude envelope correlation (oAEC) method, which minimises the effects of volume conduction and signal leakage by orthogonalisation of the tested dipole time series relative to the seed. Connectivity values were estimated for each subject and condition, and group-level contrasts between BA and RS were tested using the same cluster-based permutation procedure as applied for global functional connectivity (GFC). This approach allowed the identification of large-scale networks specifically synchronised with theta-oscillatory hub activity in V1M and the hippocampus during behavioural activity.

### Cross-frequency coupling CFC

Source localisation was performed on narrowband-filtered EEG signals spanning 0.1-100 Hz, with a resolution of 1 Hz. Seed-based functional connectivity was computed for each frequency within this range using the same connectivity metric as described for GFC. This procedure yielded a spatio–frequency map of functional connectivity with dimensions Nv × Nf, where Nv is the number of voxels and Nf the number of frequency samples. Differences between BA and RS conditions were calculated, and the resulting thresholded maps were used to identify frequency ranges of interest – specifically, low-frequency theta and high-frequency gamma bands – for subsequent CFC analysis. The goal was to determine whether CFC facilitates coupling between spatially distinct active cortical patches (29). For these selected bands, EEG signals were filtered, projected into source space, and phase–amplitude coupling (PAC) was computed within each voxel using the method of Tort et al. (30). PAC strength was quantified using the Mean Vector Length (MVL) metric (31). To correct for biases inherent in MVL estimation, values were normalised using a surrogate distribution obtained by introducing random time shifts between phase and amplitude signals over 150 iterations (31). The final PAC metric was expressed as a Z-value, representing the number of standard deviations from the surrogate mean.

## Results

### Spectral power analysis

Amplitude spectral power was compared between resting-state-like (RS) and behavioural activity (BA) conditions. A prominent increase in power was observed in a narrow spectral range from 6 to 8 Hz during BA, with a clearly identifiable peak around 7 ± 1 Hz. This theta peak was selected as a region of interest for further seed-based functional connectivity and cross-frequency coupling analyses. The grand-average power spectra across all electrodes and recordings (Fig. 1A) showed a global increase in both the theta peak and gamma-band power above 30 Hz. When spectra of individual electrodes were inspected (Fig. 1B), the theta peak was found to be most pronounced over parietal (P3/P4) and temporal (T3/T4) sites, which was consistent with the T-statistic map (Fig. 1D). In contrast, spectral power in lower frequencies spanning delta through beta bands was significantly reduced across most electrodes (Fig. 1D).

A localized increase around 1 Hz was additionally detected in temporal regions. All changes were distributed largely symmetrically across hemispheres. Representative raw EEG traces (Fig. 1E) further illustrate the differences between conditions, with BA epochs characterised by more prominent rhythmic activity compared to RS.

### Canonical band source-level spectral power and global functional connectivity

Spider plots summarise the relative magnitudes of effects across canonical frequency bands. Based on this overview, detailed source-space results are presented for the delta, beta, high-beta, gamma, and high-gamma bands in spectral power (POW; Fig. 2B, left) and for the theta, beta, high-beta, and gamma bands in global functional connectivity (GFC; Fig. 2C, left).

**Figure 2.**
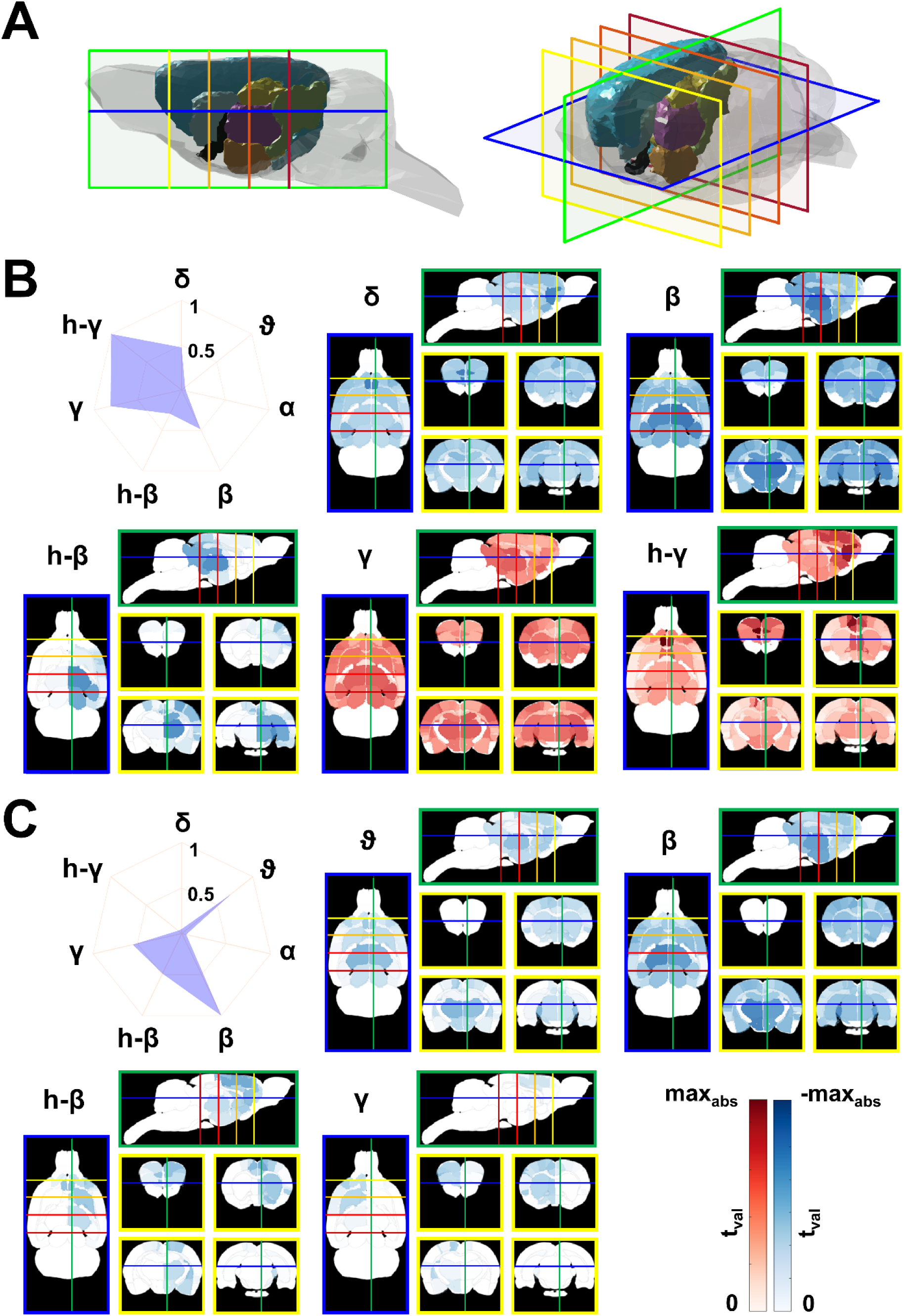
Source localisation of power and connectivity changes between behavioural activity (BA) and resting state (RS) in rats. **A)** 3D anatomical model of the rat brain indicating the positions of the displayed cutting planes: horizontal slice at -6 mm ventro-dorsally (blue), sagittal slice at +0.5 mm laterally (green), and four coronal slices at +3.5 mm (yellow), 0 mm (light orange), -3.5 mm (dark orange), and -7 mm (red) from bregma. Coloured brain structures follow the TOHOKU Rat Brain Atlas™ (28): Isocortex (light blue), Pallidum (black), Hippocampus (gold), Hypothalamus (coral), Diencephalon (purple), Midbrain (light green), and Striatum (cyan). **B)** Left: A spider plot summarizing the relative strength (normalized t-values) of effects across all frequency bands, including delta (δ, 1–4 Hz), theta (ϑ, 4–8 Hz), alpha (α, 8–12 Hz), beta (β, 12–25 Hz), high-beta (h-β, 25–30 Hz), gamma (γ, 30–40 Hz), and high gamma (h-γ, 55–100 Hz). Right: Group-level source-localised t-value maps of differences in band-limited EEG power (POW) between BA and RS conditions. Only frequency bands with significant effects are shown. Voxels with significant differences (cluster-based permutation test, p < 0.05) are shown, interpolated in Waxholm space, and mapped to corresponding anatomical regions. Warmer (red) colours indicate increased activity or connectivity during BA; cooler (blue) colours indicate decreases. **C)** Left: A spider plot summarising the relative strength (normalised t-values) of effects across all frequency bands. Right: Group-level source-localised t-value maps of differences in global functional connectivity (GFC) between BA and RS. Only frequency bands with significant effects are shown.

Within the delta band, BA was associated with a widespread reduction in power, most prominent in the medial prefrontal cortex (prelimbic/infralimbic; right t = –4.17, left t = –4.07) and anterior cingulate (Cg1; left t = –3.27, right t = –3.13). Suppression extended through insula, ventral striatum (accumbens; right t = –3.38, left t = –2.99), and sensorimotor cortex (e.g., S1Sh; left t = –3.38). Posterior cortical networks were also reduced, including parietal association, retrosplenial, and visual areas (V1; left t = –3.25). Subcortical decreases involved the diencephalon (left t = –2.64), hypothalamus (left t = –2.41), pallidum, and midbrain. No significant differences in the theta band emerged among the POWs. In the beta band, power reductions centred on diencephalon (right t = –4.37, left t = –4.06), BNST (left t = –4.16), hippocampus (right t = –4.10), and hypothalamus (left t = –4.00, right t = –3.98). These effects extended into the somatosensory cortex (S1HL; right t = –3.88) and the visual cortex (V1; left t = –3.62) as well as the retrosplenial and cingulate cortices. The high-beta band showed a right-lateralized decrease, peaking in the right diencephalon (t = –2.43) and right sensorimotor cortex (S1HL; t = –2.24), with smaller bilateral effects in the hippocampus, retrosplenial, and cingulate areas. In contrast, the gamma band exhibited a robust, widespread power increase during BA. The largest effects were found in somatosensory cortex (S1Sh; left t = 6.86), visual cortex (V1M; left t = 6.22), motor areas (M1; left t = 5.31), and hippocampal–entorhinal regions (hippocampus; right t = 6.57). Subcortical structures, including diencephalon (right t = 6.34), hypothalamus (left t = 5.85), BNST (left t = 6.59), accumbens, striatum, and septum, also showed pronounced increases. In the high-gamma band, BA was associated with strong and widespread power increases, most prominent in the medial prefrontal cortex (prelimbic/infralimbic; right t = 8.04, left t = 7.33) and anterior cingulate cortex (Cg1; left t = 7.66, right t = 6.45). Additional increases were observed in frontal association and secondary motor cortices (t ≈ 6.5), nucleus accumbens (t ≈ 6.1 right, 5.1 left), and septum (t ≈ 5.9 right; 5.7 left). Robust effects extended into orbital and insular cortices, retrosplenial regions, and subcortical structures, including hypothalamus, diencephalon, striatum, pallidum, and midbrain (t ≈ 4–5). Weaker but consistent increases were also present in hippocampal–entorhinal and posterior sensory areas. Overall, high-gamma power increases were more frontally distributed than gamma-band power increases.

In the theta band, BA led to decreased connectivity within the diencephalon (left t = –3.37), hypothalamus (left t = –2.60), ventral striatum/BNST (BNST; left t = –2.94; accumbens; right t = –2.85), and septum. Reductions were also evident in prefrontal, cingulate, motor, insula, and somatosensory regions. The beta band displayed the most extensive GFC suppression, dominated by hypothalamus (left t = –6.09) and diencephalon (left t = –5.96), with parallel decreases in BNST (left t = –5.64), striatum, amygdala, midbrain, and substantia nigra. The sensorimotor network (S1FL; left t = –5.00) and cingulate cortex (Cg1 left t = –5.04) were similarly affected, along with retrosplenial and visual cortices. In the high-beta band, connectivity decreases were smaller and somewhat right-lateralized, affecting the cingulate (Cg1; right t = –2.58), BNST, prefrontal, and motor regions, with additional reductions in the striatum, accumbens, and amygdala. Despite gamma power increases, gamma GFC was reduced, particularly in the left hemisphere across BNST (t = –2.61), frontal–cingulate–motor cortex, septum, accumbens, and striatum, indicating a decoupling of local high-frequency activity from large-scale network coherence. By contrast to POW, no significant differences were detected in the high-gamma band for GFC.

### Theta-peak (7 ± 1 Hz) source-level dynamics

Focusing on the narrow theta peak revealed a coherent pattern across metrics (Fig. 3B–C). Power in source space (POW) was strongly increased and maximally localised to the visual system and hippocampus, with peak effects in V2ML (t = 6.04), V1B (t = 5.73), V1/V1M (t = 5.70/5.70), and bilateral hippocampal formation (t = 5.72 right, t = 5.71 left). Increases extended over posterior parietal cortex (PtPD/PtPR/PtPC; t up to 5.69), retrosplenial cortex (RSGb/RSD/RSGc; t up to 5.53), lateral parietal association cortex (LPtA; t = 5.46 left, t = 4.11 right), primary somatosensory fields (S1Sh/S1Tr/S1/S1BF; t up to 5.48), temporal association/auditory cortices (TeA/AuD/Au1/AuV; t up to 5.06), and subcortically to diencephalon and hypothalamus (t = 4.93/4.90 and t = 3.95/4.10, respectively).

**Figure 3.**
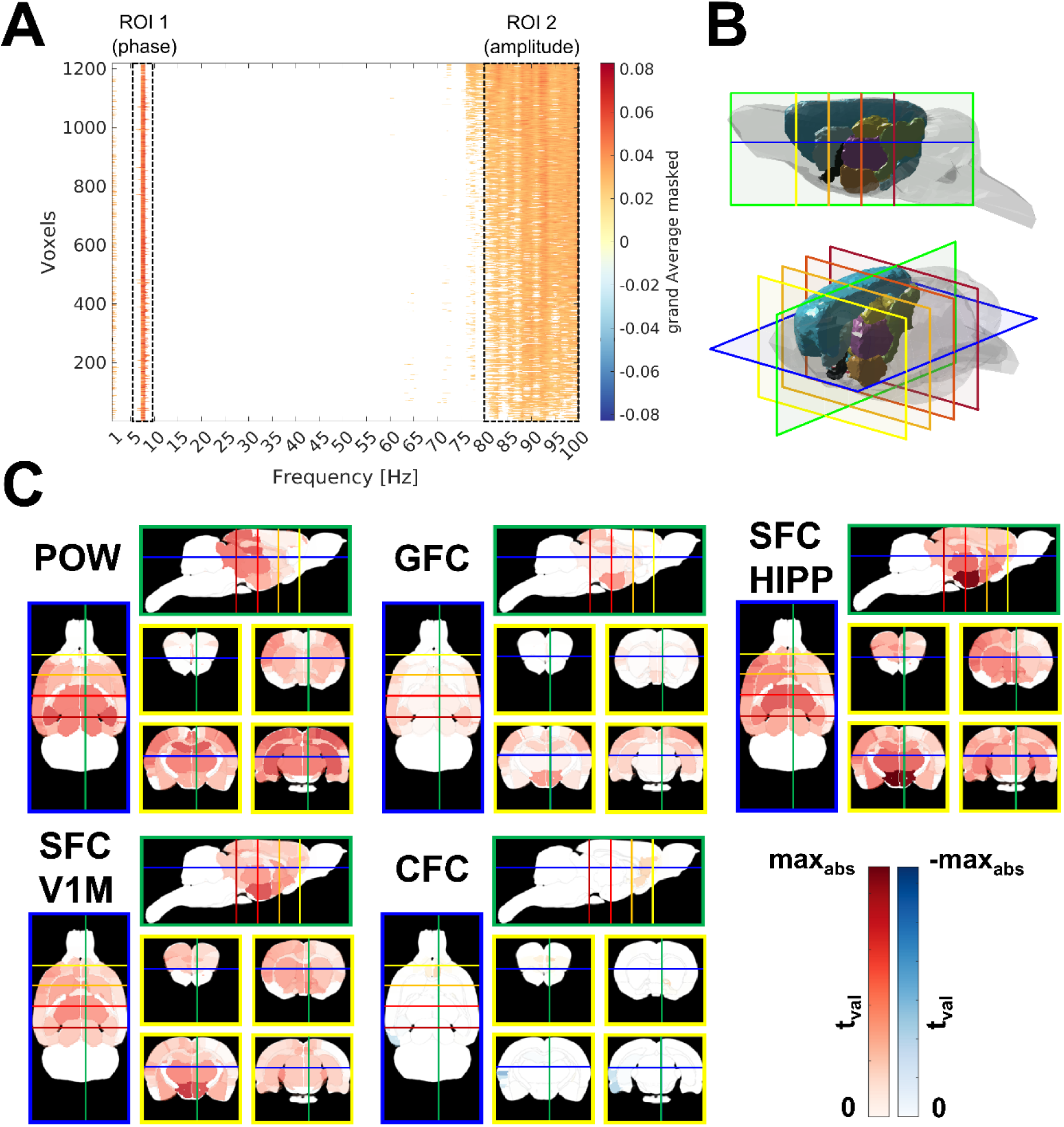
Electrophysiological correlates of behavioural activity at the theta peak frequency. **A)** Grand-averaged, statistically masked difference map of GFC (BA–RS) across frequencies (1–100 Hz) and all voxels in Waxholm space. Highlighted regions of interest (ROI 1: 6–8 Hz for phase, ROI 2: 80–100 Hz for amplitude) were used to define phase and amplitude carriers in subsequent CFC analysis. Only the differences above the third quartile are shown here (differences below this threshold are masked out). **B)** 3D anatomical model of the rat brain indicating the positions of the displayed cutting planes: horizontal slice at −6 mm dorsoventrally (blue), sagittal slice at +0.5 mm laterally (green), and four coronal slices at +3.5 mm (yellow), 0 mm (light orange), −3.5 mm (dark orange), and −7 mm (red) from bregma. **C)** Source-localised statistical t-maps showing significant changes between behavioural activity (BA) and resting state (RS) in: – POW: power in source space, – GFC: global functional connectivity, – SFC: seed-based functional connectivity from hippocampus (HIPP) and right V1M (primary visual cortex), and – CFC: cross-frequency coupling between theta (phase) and gamma (amplitude) frequencies. Maps show only significant areas (cluster-based permutation test, p < 0.05), colour-coded by t-value (red indicates an increase in BA, blue indicates a decrease).

Global functional connectivity (GFC) in the theta peak was increased and centred on posterior sensory–parietal hubs and hypothalamus. The largest effects were observed in the right hypothalamus (t = 3.47) and left hypothalamus (t = 3.36), followed by right V1B (t = 3.42), right MPtA (t = 3.37), right V2L/V2ML/V2MM (t up to 3.33), bilateral posterior parietal subfields (PtPD/PtPR/PtPC; t up to 3.26), and primary somatosensory regions (S1/S1Tr/S1BF/S1Sh; t up to 3.10). Additional increases were present in retrosplenial cortex (RSGc/RSD; t ≈ 2.47–2.20), temporal/auditory cortices (TeA/AuD/Au1/AuV; t ≈ 2.83–2.45), hippocampal formation (t = 2.54 left, t = 2.54 right), and preoptic/BNST regions (t = 2.12–1.76).

To test whether the posterior visual cortex or hippocampus acts as a hub at this frequency, seed-based functional connectivity (SFC) was computed using right V1M and the whole hippocampus as the seed. For V1M, SFC increases were strongest in subcortical integrative nodes: hypothalamus (t = 5.68 left, t = 5.13 right), BNST (t = 4.76 left, t = 4.10 right), preoptic area (t = 4.32 right, t = 3.72 left), and diencephalon (t = 4.22 left, t = 3.92 right). Robust effects also involved insular (GI/DI; t up to 4.01), cingulate (Cg1/Cg2; t up to 3.60), retrosplenial (RSGc/RSGb/RSD; t up to 3.91), basal ganglia/striatal territories (striatum/pallidum/substantia nigra; t up to 3.72), septum (t ≈ 3.88–3.88), anterior commissure (t = 3.63 left, t = 3.63 right), and several primary somatosensory subfields (S1DZO/S1J/S1FL/S1ULp/S1HL; t up to 4.01). Hippocampus and entorhinal cortex were also engaged (t ≈ 2.74–3.13). For the hippocampus, SFC increases were the strongest.

Finally, cross-frequency coupling (CFC) using theta-phase (6–9 Hz) and within-high-gamma-amplitude (80–100 Hz) revealed focal increases and predominantly left-lateralized decreases (Fig. 3A, B). PAC increased in medial prefrontal cortex—prelimbic/infralimbic (t = 1.18 left, 0.80 right)—and in right nucleus accumbens (t = 1.16) and right frontal association cortex (t = 1.13). Decreases were strongest in the left insular cortex (GI+DI; t = –1.92), left dorsal-intermediate entorhinal cortex (t = –1.55), left perirhinal and ectorhinal cortices (t = –1.37 and –1.35), and left ventral-auditory cortex (t = –1.38), with additional modest reductions across left medial/ventral entorhinal and temporal association areas.

## Discussion

Behavioural activity in freely moving rats is associated with a characteristic, frequency-specific reorganisation of EEG dynamics that was consistent across power (POW), global functional connectivity (GFC), seed-based connectivity (SFC), and cross-frequency coupling (CFC). We first identified relative effect magnitudes across bands, then focused source-space analyses on those bands showing robust effects.

The most salient spectral feature of active behaviour was a narrowband theta peak at ∼7 ± 1 Hz. Source localisation placed this increase predominantly in posterior sensory cortices (visual, somatosensory, auditory) and hippocampal formation, with little or no engagement of the primary motor cortex. Such a topography argues against movement artefact and instead points to recruitment of sensory, associative, and mnemonic circuitry during exploration. The association between voluntary movement and theta oscillations in the 6–8 Hz range is well-established (32,34,35). Theta power and phase organise hippocampal and parahippocampal firing during navigation and decision-making, scale with locomotor speed, and support sensorimotor integration and spatial memory (36–40). Theta also accompanies intentional acts beyond locomotion, including frontal theta during consummatory behaviours such as licking (41), underscoring its broad role in coordinating action with sensory feedback. Our localisation to retrosplenial and posterior parietal fields dovetails with their established contributions to visuospatial orientation and memory-guided locomotion (35,38) and is consistent with our earlier electrode-level indications of a parietal maximum for the ∼7 Hz peak (4,20).

Gamma power increased widely across the cortex and subcortex during behavioural activity, with prominent effects in sensory and motor neocortex, hippocampus–entorhinal areas, basal ganglia, and diencephalon. Gamma oscillations index local circuit engagement and are enhanced by arousal, sensorimotor processing, and active environmental sampling. In rodents, running amplifies gamma activity in the visual cortex, thereby improving visual processing during movement. Additionally, gamma activity in motor/somatosensory regions accompanies movement initiation and proprioceptive feedback (42–44). Functionally, a global gamma upshift implies that the brain enters a computation-rich regime in which fast local assemblies dominate information transmission and sensory–motor integration (43). Gamma often couples to theta such that theta phase modulates gamma amplitude, a mechanism proposed to organise fine-scale (gamma) content within theta cycles during active cognition (45–47). The parallel upregulation of theta and gamma power in our data thus aligns with an “activated” cortical state, in which posterior sensory–hippocampal networks and subcortical integrators cooperate to support the exploration and guidance of movement.

Countervailing decreases in lower frequencies were equally systematic. Delta power (1–4 Hz) fell broadly, with maxima in medial prefrontal and anterior cingulate regions and extensions into insula, ventral striatum, and posterior cortices. Delta is a hallmark of drowsiness, slow-wave sleep, and pharmacological sedation; it diminishes with arousal and behavioural activation (48). Our delta suppression, therefore, may reflect a state transition from an idling, synchronous mode to an alert, desynchronized mode characteristic of active wakefulness.

Beta-band power (12–25 Hz) and high-beta power (25–30 Hz) also declined, with the strongest beta reductions observed in the diencephalon, hypothalamus, basal ganglia, hippocampus, and sensorimotor/visual cortices, and a right-lateralized pattern in high-beta power. Movement-related beta desynchronization is a robust phenomenon in both human and animal studies, indexing release from a status quo motor set and enabling movement initiation and updating (49–53). The involvement of basal ganglia and diencephalic structures in our maps is consistent with beta’s generation within cortico-basal ganglia–thalamic loops and with dopaminergic modulation of these rhythms. Pathologically exaggerated beta in Parkinson’s disease reflects impaired dopamine transmission, whereas dopaminergic stimulation tends to reduce beta synchrony (54,55). Pharmacological evidence in rodents likewise shows that stimulants decreasing beta can accompany hyperlocomotion, while dopamine antagonism can elevate beta (56–59). Mechanistically, the beta decrease during movement plausibly arises from increased inhibitory drive and desynchronization within GABAergic interneuron networks and basal ganglia output nuclei, thereby facilitating flexible motor output (60–62). The right-dominant high-beta suppression we observed resembles lateralized beta desynchronization reported with asymmetric motor preparation and may reflect hemispheric specialisation or subtle behavioural asymmetries (63).

Connectivity analyses revealed a complementary reorganisation at the network level. Using orthogonalized amplitude-envelope correlations to index GFC, we found predominantly reduced long-range synchronisation during behavioural activity in broad theta (4–8 Hz), beta, high-beta, and gamma ranges. Suppression was strongest in thalamic/hypothalamic compartments, the BNST/ventral striatum, the amygdala, midbrain, and substantia nigra, and extended to the cingulate, retrosplenial, visual, and motor cortices. This decoupling, despite concurrent power increases in higher bands, suggests a shift toward segregated, locally driven processing during active states: fast rhythms amplify within regions, while slow inter-areal coherence recedes. The pattern is expected for gamma, whose spatial coherence is typically local, and accords with the known breakdown of resting-state beta/theta coherence when animals initiate movement (64,65). It also helps reconcile discrepancies between natural behaviour and pharmacological hyperlocomotion: drug-induced states (e.g., NMDA antagonists, psychostimulants) can introduce additional, sometimes aberrant coherences, whereas voluntary movement alone primarily diminishes slow-band global coupling (4,33). These findings reinforce the importance of controlling for locomotor state in pharmaco-EEG, because active versus inactive periods differ markedly in baseline connectivity, potentially obscuring or mimicking drug effects (33).

An important exception to the overall GFC reduction emerged at the narrow theta peak (∼7 Hz), where connectivity increased among posterior sensory–parietal hubs and key subcortical nodes. Theta-peak GFC rose in primary/secondary visual cortex, posterior parietal and retrosplenial areas, primary somatosensory fields, and in hypothalamus/diencephalon. This focal enhancement supports the view that the locomotor theta rhythm coordinates a task-specific communication backbone linking perception, spatial mapping, and state control. The hypothalamus and adjacent diencephalic structures participate in locomotor drive, head-direction, and arousal systems; their increased theta coupling with posterior cortex points to tight integration of environmental sampling with internal state and motor set during exploration (36,66,67). In humans, analogous theta-range cortico-hippocampal/parietal couplings emerge during active navigation, suggesting a conserved role for theta in binding distributed sensorimotor computations into a coherent navigation loop.

Seed-based analysis centred on the right V1M at 7 Hz further clarified the architecture of this loop. During behavioural activity, V1’s theta phase/envelope synchronisation strengthened with hypothalamus, BNST, preoptic area, and diencephalon, as well as with insula (GI/DI), cingulate (Cg1/Cg2), retrosplenial subfields, basal ganglia (striatum, pallidum, substantia nigra), septum, and commissural pathways; hippocampus and entorhinal cortex were also engaged. This pattern suggests that sensory cortical oscillations are temporally aligned with subcortical integrators that govern arousal, motivation, and movement vigour. BNST and preoptic–hypothalamic regions regulate contextual anxiety, reward seeking, thermoregulation, and autonomic set; their coupling to the visual cortex at the theta peak suggests that salient visual cues during exploration are rapidly routed to systems that shape locomotor strategy and internal milieu. The concomitant engagement of retrosplenial/posterior parietal areas, crucial for route computation and spatial memory, completes a plausible circuit by which theta aligns environmental sampling, internal state regulation, and motor output.

Crucially, we computed cross-frequency coupling and found that behavioural activity reconfigured theta–gamma phase–amplitude coupling (PAC) in a region-specific manner. PAC increased in medial prefrontal cortex (prelimbic/infralimbic) and in right nucleus accumbens and right frontal association cortex, but decreased predominantly in the left hemisphere across insular cortex (GI+DI), dorsal-intermediate entorhinal cortex, perirhinal/ectorhinal cortices, and ventral auditory cortex. Theta–gamma PAC has been proposed to underpin multiscale information processing—embedding gamma-encoded content within theta cycles to coordinate encoding, working memory, and decision-making (45–47,68–70). The observed PAC enhancement in the mPFC and accumbens aligns with their roles in action selection, reward prediction, and top–down control during exploration, where aligning local gamma bursts with theta phase may optimise gating and routing of task-relevant information (71). Conversely, PAC reductions in insular and entorhinal/perirhinal territories suggest a dynamic reprioritisation: during active outward-oriented behaviour, interoceptive and certain mnemonic gateway computations may be less tightly bound to theta timing, while posterior sensory–parietal and frontostriatal loops take precedence. Together with the theta-peak GFC/SFC results, these PAC effects indicate that behavioural activity does not merely scale oscillations but retunes which cross-frequency channels are used by which circuits, emphasising frontostriatal control and posterior sensory–hypothalamic integration while down-weighting others.

Several limitations should be acknowledged. First, although the rat EEG source-localisation framework enables area-level inferences (1), resolution for deep structures remains inherently constrained by the EEG inverse problem and by the distribution of surface sensors (72). This is particularly relevant to our findings in the hypothalamic, BNST, and midbrain; convergent evidence from depth electrophysiology would therefore be highly valuable. Second, spontaneous behaviour is heterogeneous; while our BA/RS labelling minimised gross confounds, a finer parsing of behavioural sub-states (e.g., gait, rearing, grooming) might reveal more specific oscillatory motifs. Third, an additional limitation concerns the electrode system. Although our 14-channel array provides above-standard coverage for rodent EEG and samples the most relevant cortical regions, it does not uniformly cover the cortical surface. A denser, more homogeneous electrode layout would likely increase the spatial precision of source localisation and refine the anatomical interpretation of the results.

## Conclusion

In conclusion, behavioural activity reorganises rat brain dynamics along clear spectral and network axes. Theta (∼7 Hz) and gamma power increases—localised to posterior sensory cortices, hippocampus, and subcortical integrators—coexist with widespread delta and beta power decreases, including a right-dominant high-beta desynchronization in sensorimotor and diencephalic territories. At the network level, GFC diminishes across broad theta, beta, high-beta, and gamma bands, indicating reduced long-range coherence during movement. In contrast, the narrow theta peak supports stronger, task-specific coupling among posterior sensory–parietal hubs and the hypothalamus. Seed-based analyses from the visual cortex highlight theta synchronisation with hypothalamic/BNST and basal ganglia nodes, and CFC reveals a redistribution of theta–gamma coupling that emphasises medial prefrontal and accumbens circuits while attenuating coupling in insular and entorhinal/perirhinal regions. Together, these findings outline a coherent physiological picture: in active wakefulness, the rat brain shifts toward a locally amplified, selectively coupled, theta-anchored regime optimised for visuospatial navigation, action selection, and state control. Beyond providing a behaviourally grounded baseline for pharmaco-EEG (4,33), this framework should help parse how neuromodulators and candidate therapeutics reshape the interplay of power, connectivity, and cross-frequency coupling across sensory, limbic, and motor circuits.

## Acknowledgment

This work was supported by the Science Foundation of Charles University in Prague (project no.: 385221), Czech Science Foundation (project no.: 23-07578K), Ministry of the Interior of the Czech Republic (project VK01010212), Long-term conceptual development of research organization (RVO 00023752), and program INTER-EXCELLENCE (subprogram INTER-ACTION LUAIZ24146), ERDF-Project Brain dynamics, No. CZ.02.01.01/00/22_008/0004643, project VVI CZECRIN (LM2023049), PsyPal project from Horizon Europe (grant no. 101137378, HORIZON-HLTH-2023-DISEASE-03-01) and Charles University research program Cooperatio-Neurosciences and private funds obtained via PSYRES, Psychedelic Research Foundation (https://psyresfoundation.eu).

## Conflict of interest

Tomáš Páleníček declares to have shares in „Psyon s.r.o.,“ and in “Společnost pro podporu neurovědního výzkumu s.r.o.” He founded „PSYRES – Psychedelic Research Foundation“. TP also reports consulting fees from GH Research and CB21-Pharma outside the submitted work. TP is involved in Compass Pathways trials with psilocybin and the MAPS clinical trial with MDMA outside the submitted work.

## Notes

### Competing Interest Statement

Tomas Palenicek declares to have shares in Psyon s.r.o., and in Spolecnost pro podporu neurovedniho vyzkumu s.r.o. He founded PSYRES, Psychedelic Research Foundation. TP also reports consulting fees from GH Research and CB21-Pharma outside the submitted work. TP is involved in Compass Pathways trials with psilocybin and the MAPS clinical trial with MDMA outside the submitted work.

